# Sample delivery using capillary and delivery medium for serial crystallography

**DOI:** 10.1101/696138

**Authors:** Ki Hyun Nam

## Abstract

Serial crystallography (SX) is an innovative technology in structural biology that enables the visualization of molecular dynamics of macromolecules at room temperature. SX experiments always require a considerable amount of effort to deliver a crystal sample to the X-ray interaction point continuously and reliably. Here, a sample delivery method using a capillary and a delivery medium is introduced. The crystals embedded in the delivery medium can pass through the capillary tube, which is aligned with the X-ray beam, at very low flow rates without requiring elaborate delivery techniques and drastically reducing sample consumption. This simple but highly efficient sample delivery method can allow researchers to deliver crystals precisely to X-rays in SX experiments.

## Introduction

Traditional X-ray crystallography has contributed greatly to biology, medicine, and industrial fields;^1–3^ however, this method has certain experimental limitations, including radiation damage, cryogenic temperature, and static structural information.^4,5^ This technique often produces biologically less relevant crystal structures,^6,7^ but its limitations can be overcome by the serial crystallography (SX) technique. An SX experiment using X-ray free electron laser (XFEL) or a synchrotron X-ray source allows radiation damage-free or low-dose data collection of macromolecules, respectively.^8–10^ This technique determines the room-temperature structure of crystals; moreover, time-resolved molecular dynamics in the range of femtoseconds to a few seconds can be determined through a pump-probe study.^10,11^ Therefore, SX can help develop new approaches to structural biology beyond the experimental limitations of traditional X-ray crystallography.

SX experiments require crystal samples to be delivered to the X-ray interaction point in a steady and stable manner.^12^ Various sample delivery techniques such as injector-based methods,^13,14^ fixed-target scanning,^15,16^ electrospinning,^17^ and methods employing microfluidic devices^18^ have been developed, and their application has been successfully demonstrated in SX experiments. However, most of these methods require precisely fabricated sample delivery devices and are time consuming and expensive to perform. In addition, considerable amounts of effort and skill are required to operate the delivery system precisely during data acquisition, as opposed to easily mounting single crystals in conventional X-ray crystallography using a synchrotron.

Among these techniques, the injector-based sample delivery method is particularly popular and is widely applied in the SX experiments.^13,14,19^ As a liquid jet sample injector can maintain a stable injection stream at high flow rates,^13^ the typical sample consumption is very high for XFEL facilities with a low repeat rate or those using a synchrotron.^14,20^ Meanwhile, sample delivery using an injector with a viscous delivery medium can drastically reduce the sample consumption at low flow rates.^19,21^ However, the viscosity of the delivery medium varies depending on the crystallization solution and experimental environment; therefore, considerable effort is still required to maintain a stable injection stream. In contrast, a method of delivering a crystal suspension using a capillary was introduced previously.^22^ Although this approach has been successful in delivering crystal samples, there is a drawback in that the sample consumption is high owing to the high flow rate of 2.5 μl/min. In addition, this method involves the problem of crystals sinking in the capillary due to gravity at low flow rates. Therefore, a straightforward and universal sample delivery system should be established to ensure stable delivery of crystal samples to X-rays that can minimize sample consumption and does not require professional operating skills.

This paper proposes a straightforward sample delivery method using a capillary and a delivery medium for SX. The capillary fixes the path of the delivery medium such that crystals are delivered to the X-ray interaction point perfectly without the need for precise operation. The delivery medium not only prevents crystals from sinking into the capillary but also allows delivery in extremely low flow rates, which considerably reduces sample consumption. This method was successfully applied to the serial millisecond crystallography (SMX) experiment and could determine the room-temperature crystal structure of lysozyme and glucose isomerase at 1.85 Å and 1.80 Å resolution, respectively, using less than 200 μg of protein crystals.

## Results and discussion

A capillary is an X-ray transparent material that generates low background scattering, which is negligible in data processing; thus, it has been often used to mount crystals in traditional X-ray crystallographic studies.^23^ When the center of the capillary is aligned with the X-ray beam path, the sample injected into the capillary only follows the predetermined path. As a result, all samples follow the X-ray beam path and sophisticated delivery operating skills are not required. However, when low flow rates are used to reduce sample consumption in this state, the crystal samples can sink to the bottom of the capillary due to gravity. In such a state, when the crystal samples at the bottom of the capillary are physically moved to the X-ray position by a syringe plunger, a large number of crystals are simultaneously exposed to X-rays, causing data processing problems owing to multi-hit patterns; this may also lead to clogging, causing the capillary to break or become damaged. To solve this problem, a delivery medium was used in this experiment to ensure that the crystals do not settle by gravity even at low flow rates. In general, when delivering crystals using a viscous delivery medium in SX experiments, it is essential to determine the optimal injection condition required for a stable injection stream based on the crystallization solution and experimental environment. In contrast, the delivery medium used in this experiment intended to prevent crystals from sinking in the capillary due to gravity; thus, pre-experiments to determine a stable injection stream condition were not required. As the crystal samples embedded in the delivery material pass through the fixed capillary, the sample can be delivered at extremely low flow rates, which enables sample consumption to be drastically reduced compared to conventional methods.

To demonstrate sample delivery using a capillary and a delivery medium, SMX experiment was performed at a synchrotron source. A quartz capillary with an inner diameter of 200 μm was connected to the tip of a commercial syringe needle. The tip of the needle was firmly attached to the wide inlet of the capillary so that the delivery medium did not leak; the capillary and the syringe needle were fixed with polyimide tape (**Figure 1A**). The sealed end of the capillary was opened using a cutter so that delivery medium containing the crystals could pass continuously (**Figure 1A**). In order to prove that the delivery medium used in this method only serves to prevent crystals from sinking in the capillary, agarose and gelatin materials were used as a delivery medium. Agarose is known to be an effective sample delivery medium in vacuum environments and no methods have been reported to provide a stable injection stream at atmosphere at room temperature.^24^ Gelatin has been reported as an unsuitable delivery medium.^24^ A crystal suspension of lysozyme and glucose isomerase, as a model sample, was mixed with the agarose and gelatin using a dual syringe-setup. The syringe containing the crystals embedded in the delivery medium was connected to the capillary-associated needle (Figure 1), which was installed into the syringe pump. After alignment of the capillary into the X-ray beam pass using a manual stage, the crystals embedded in the delivery medium were extruded from the syringe by the driving force of the syringe pump. The sample was delivered at a flow rate of 100 nl/min in a capillary with a diameter of 400 μm.

**Figure 1.**
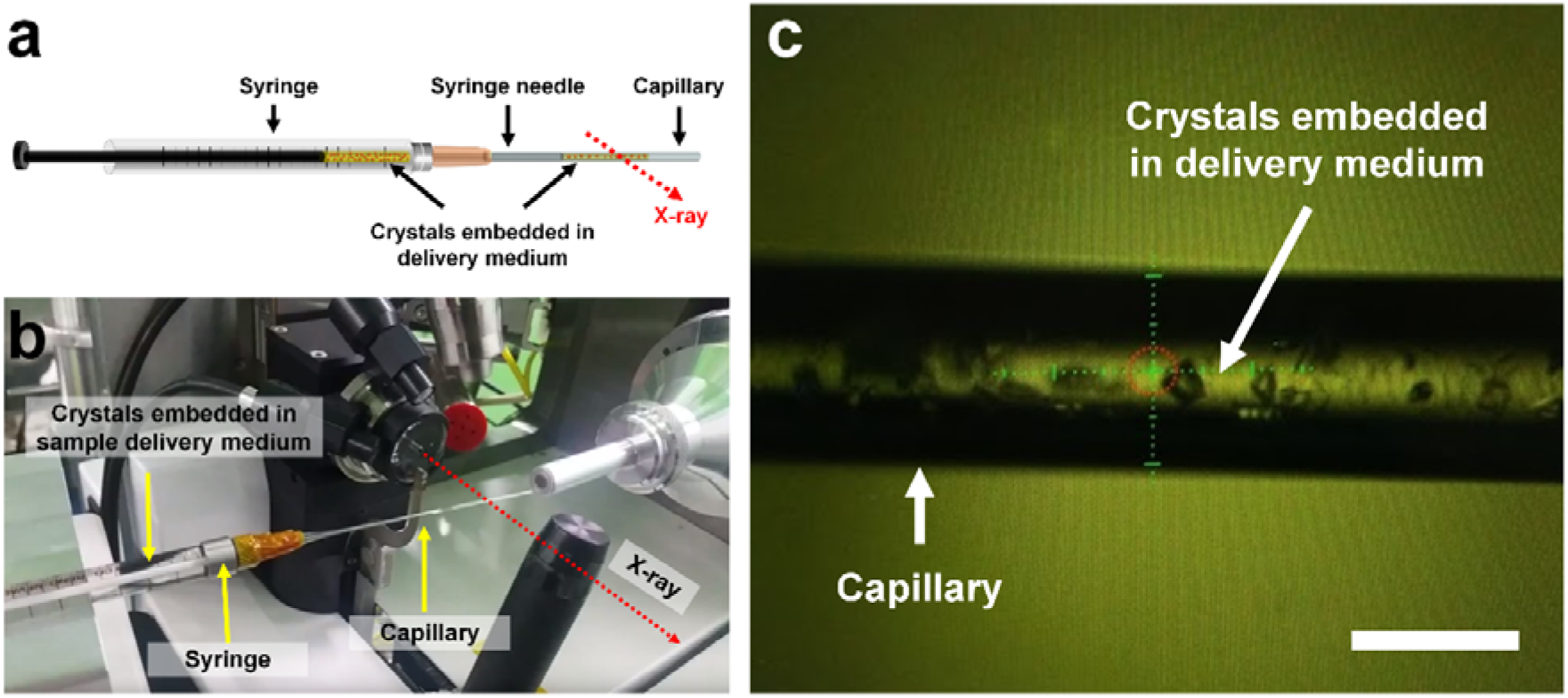
Sample delivery using a capillary and a viscus medium. (a) Schematic of the capillary tube connected to the syringe. (b) Experimental setup of sample delivery using the capillary and viscus medium. Capillary was aligned to the X-ray beam path. Crystals embedded in the delivery medium were extruded from the syringe by the driving force from the syringe pump. (c) Snapshot of the capillary used to deliver lysozyme crystals embedded in an agarose delivery medium. Scale bar indicates 400 μm.

The X-ray energy used was 12.659 keV with a flux of 1.3 x 10^12^ photons/s. The X-ray beam, which was focused using a K-B mirror, had a size of 4.1 μm (vertical) × 8.5 μm (horizontal) (full width at half maximum) at the sample position. Crystals were exposed to the X-ray for 100 ms, and data were collected at 25 °C. A total of 48000 diffraction data points were each collected for lysozyme and glucose isomerase. For the lysozyme, 34378 indexed images were obtained and processed up to 1.85 Å (Supplementary Figure S1); the overall SNR, completeness, CC, and R_split_ were 6.33, 100, 0.9950, and 6.58, respectively. For the glucose isomerase, 43197 indexed images were obtained and processed up to 1.70 Å (Supplementary Figure S2); the overall signal-to-noise ratio (SNR), completeness, pearson correlation coefficient (CC), and Rsplit were 5.15, 100, 0.9819, and 12.41, respectively. In the diffraction images collected, no background scattering was observed owing to the capillary that could affect data processing. The R_work_/R_free_ of the lysozyme and glucose isomerase structure were 17.33/22.44 and 19.26/22.36, respectively. The glucose isomerase and lysozyme showed a very clear electron density map for Lys19-Leu147 and Tyr3-Arg387, respectively (Figure 2). In these structures, no significant radiation damage was observed at the disulfide or metal binding sites (Supplementary Figure S3).

**Figure 2.**
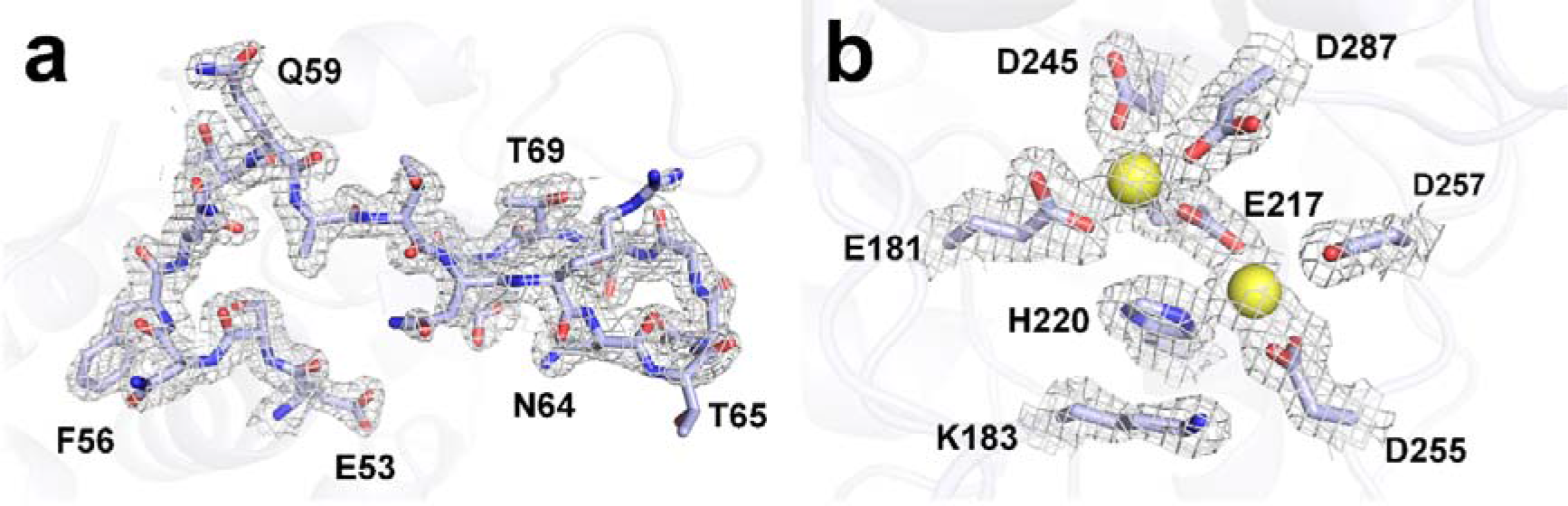
2Fo-Fc electron density map (grey mesh, 1.5σ) of (a) lysozyme delivered in agarose and (b) glucose isomerase delivered in gelatin.

In summary, a sample delivery system using a capillary and viscous medium has been introduced and successfully used to determine the structure of glucose isomerase and lysozyme at room temperature with agarose and gelatin. This approach can enable researchers to deliver samples in a simple and precise manner in SX experiments and can be applied to deliver various samples from macromolecules to small molecules. The selected delivery medium does not have any physical or chemical effects on the crystal samples. Hydrogel-like materials can be effectively used in this method as they generate no or very weak background scattering. This simple method encourages researchers to further investigate SX experiments at room temperature.

## Materials and methods

### Preparation of capillary-connected syringe needle

Quartz capillary tubes with a diameter of 400 μm were purchased from Hampton Research (HR6-134). The ends of syringe needles were tightly adhered to the large opening of the quartz capillary and then fixed with polyimide tape.

### Sample preparation

Lysozyme crystals from a chicken white egg and glucose isomerase *Streptomyces rubiginosus* were prepared as previously reported.^16^ The sizes of the lysozyme and glucose isomerase crystals used in the experiments were 30–50 μm and <60 μm, respectively. A crystal suspension of lysozyme (20 μl) and glucose isomerase (20 μl) was embedded in 5% (w/v) agarose (30 μl) and 5% (w/v) gelatin (30 μl), respectively, by mechanical mixing in a dual syringe setup.^19^ Crystals embedded in the delivery medium were transferred into a 100 μl syringe; this syringe was connected to the capillary-associated needle.

### Data collection

SMX was performed at an 11C microbeamline at PLS-II.^25^ The photon energy and flux were 12.659 keV and 1.3 x 10^12^ photons/s, respectively. The crystal sample was exposed to the X-ray for 100 ms. The temperature and humidity of the experimental hutch were 25 °C and 20%, respectively. The capillary-connected syringe was installed into the Fusion Touch 100 syringe pump (CHEMYX). The capillary was aligned to the X-ray beam path. The crystal sample embedded in the delivery medium was extruded from the syringe by pushing the plunger using a driving motor.^26^ The sample flow rate was 200–300 nl/min. The diffraction patterns were recorded on Pilatus 6M with a 10 Hz readout.

### Structure determination

The diffraction patterns were filtered and processed using the Cheetah^27^ and CrystFEL^28^ programs, respectively. Molecular replacement was performed using Phaser-MR^29^ with the crystal structure of lysozyme (PDB code 6IG6)^21^ or glucose isomerase (5ZYE)^30^ as the search modes. The model was established and its structure was refined using the COOT^31^ and PHENIX^32^ programs, respectively. The geometry was analyzed using MolProbity^33^. All figures were generated by PyMOL (https://pymol.org/). Data collection and refinement statistics are summarized in Supplementary Table 1.

### Accession codes

Coordinates and structure factors have been deposited under accession codes 6KD1 (lysozyme delivered in agarose) and 6KD2 (glucose isomerase delivered in gelatin) in Protein Data Bank.

## Acknowledgments

I thank the beamline staff at 11C beamline at Pohang Accelerator Laboratory for their assistance with data collection. The authors thank Global Science experimental Data hub Center (GSDC) at Korea Institute of Science and Technology Information (KISTI) for computational support. This work supported by a Korea University Grant. This work was funded by the National Research Foundation of Korea (NRF-2017M3A9F6029736).

## AUTHOR CONTRIBUTIONS

Ki Hyun Nam performed all experiment and wrote the manuscript.

---

**Supplementary Table 1.**
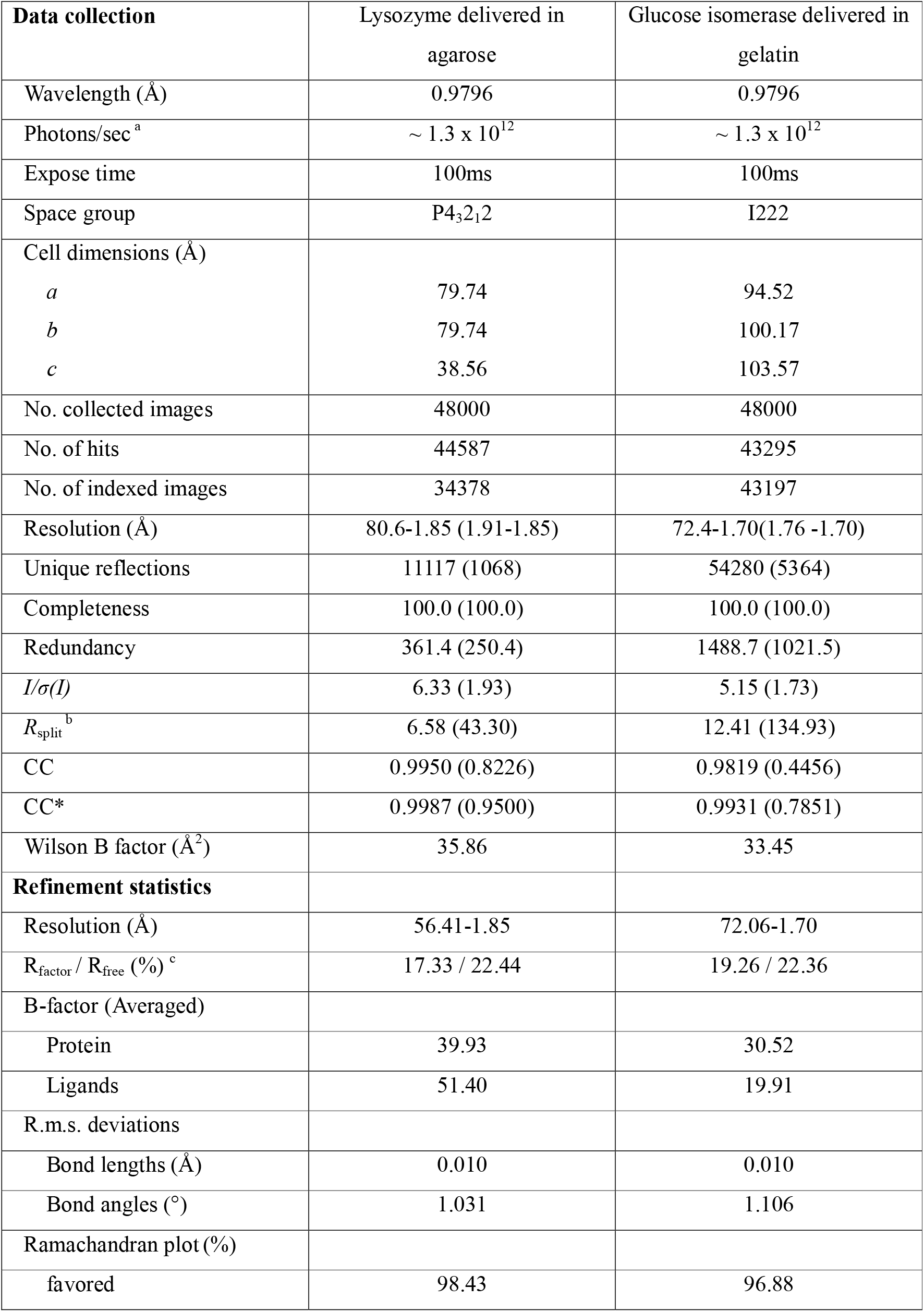
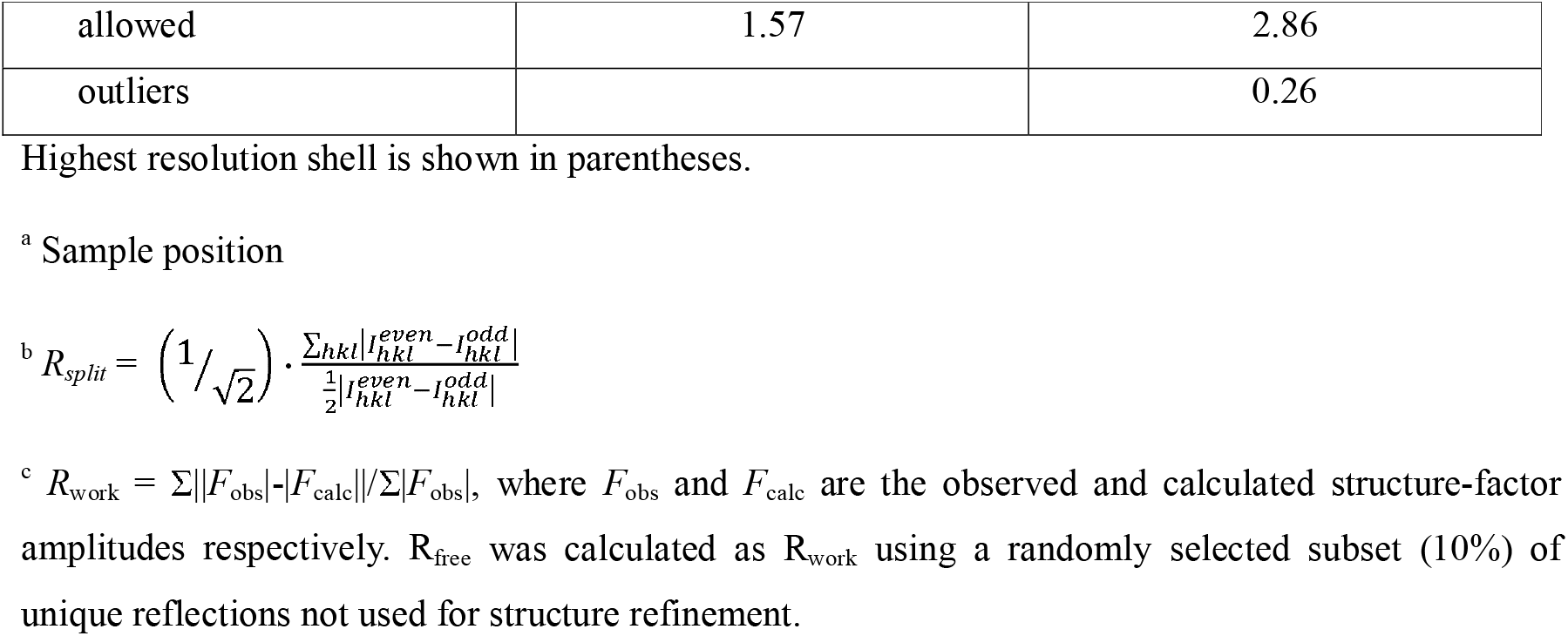
Data collection and refinement statistics.

| **Data collection** | Lysozyme delivered in<br>agarose | Glucose isomerase delivered in<br>gelatin |
| --- | --- | --- |
| Wavelength (Å) | 0.9796 | 0.9796 |
| Photons/sec<sup>a</sup> | $\sim 1.3 \times 10^{12}$ | $\sim 1.3 \times 10^{12}$ |
| Expose time | 100ms | 100ms |
| Space group | P4<sub>3</sub>2<sub>1</sub>2 | I222 |
| Cell dimensions (Å) |  |  |
| *a* | 79.74 | 94.52 |
| *b* | 79.74 | 100.17 |
| *c* | 38.56 | 103.57 |
| No. collected images | 48000 | 48000 |
| No. of hits | 44587 | 43295 |
| No. of indexed images | 34378 | 43197 |
| Resolution (Å) | 80.6-1.85 (1.91-1.85) | 72.4-1.70(1.76 -1.70) |
| Unique reflections | 11117 (1068) | 54280 (5364) |
| Completeness | 100.0 (100.0) | 100.0 (100.0) |
| Redundancy | 361.4 (250.4) | 1488.7 (1021.5) |
| *I*/$\sigma$(*I*) | 6.33 (1.93) | 5.15 (1.73) |
| *R*<sub>split</sub><sup>b</sup> | 6.58 (43.30) | 12.41 (134.93) |
| CC | 0.9950 (0.8226) | 0.9819 (0.4456) |
| CC* | 0.9987 (0.9500) | 0.9931 (0.7851) |
| Wilson B factor (Å<sup>2</sup>) | 35.86 | 33.45 |
| **Refinement statistics** |  |  |
| Resolution (Å) | 56.41-1.85 | 72.06-1.70 |
| *R*<sub>factor</sub> / *R*<sub>free</sub> (%)<sup>c</sup> | 17.33 / 22.44 | 19.26 / 22.36 |
| B-factor (Averaged) |  |  |
| Protein | 39.93 | 30.52 |
| Ligands | 51.40 | 19.91 |
| R.m.s. deviations |  |  |
| Bond lengths (Å) | 0.010 | 0.010 |
| Bond angles (°) | 1.031 | 1.106 |
| Ramachandran plot (%) |  |  |
| favored | 98.43 | 96.88 |
| allowed | 1.57 | 2.86 |
| outliers |  | 0.26 |

Highest resolution shell is shown in parentheses.
<sup>a</sup> Sample position
<sup>c</sup> $R_{work} = \sum ||F_{obs}| - |F_{calc}|| / \sum |F_{obs}|$ , where $F_{obs}$ and $F_{calc}$ are the observed and calculated structure-factor amplitudes respectively. $R_{free}$ was calculated as $R_{work}$ using a randomly selected subset (10%) of unique reflections not used for structure refinement.

## Supplementary Figures

**Supplementary Figure S1.**
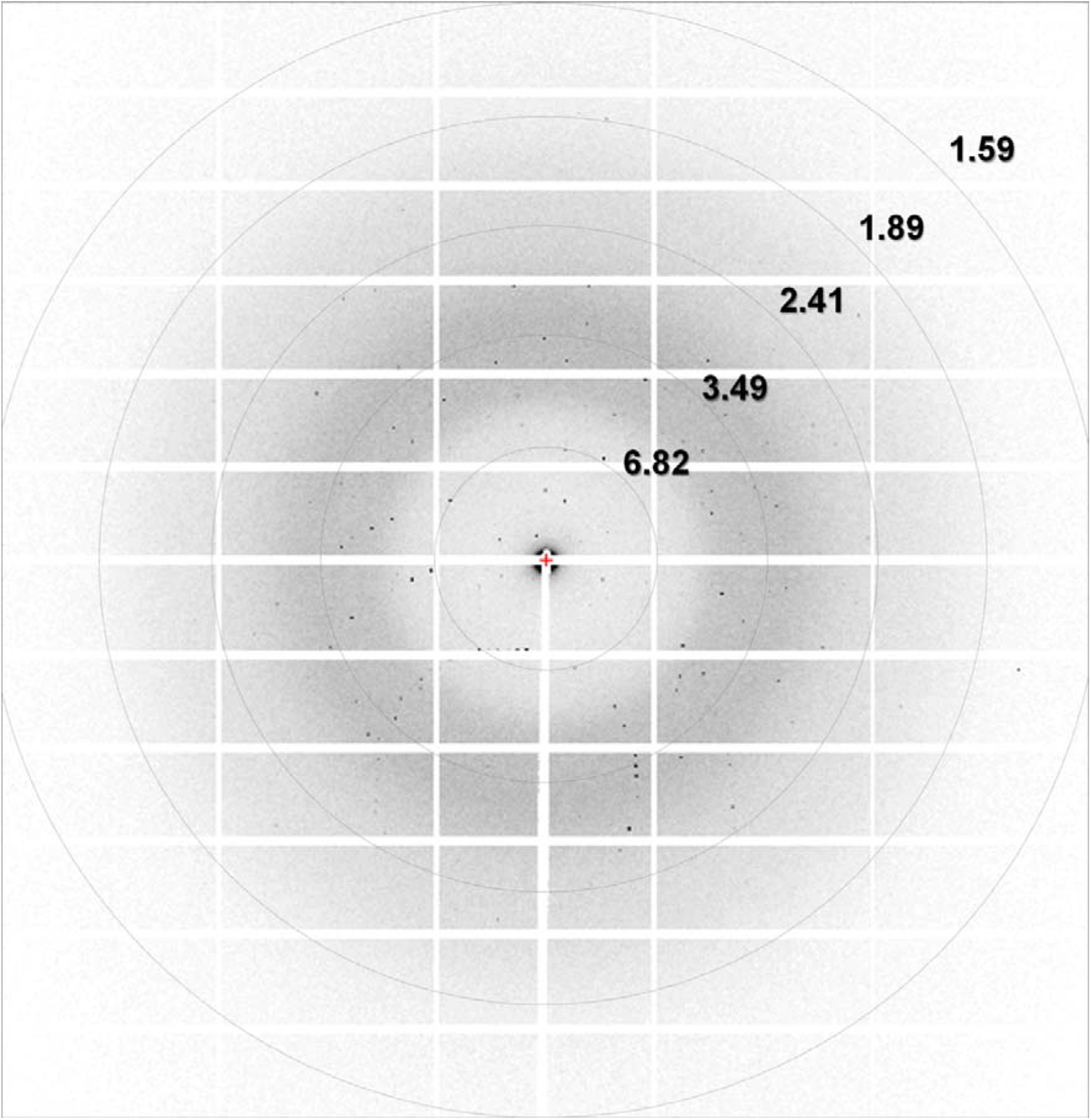
Diffraction pattern of lysozyme delivered in agarose.

**Supplementary Figure S2.**
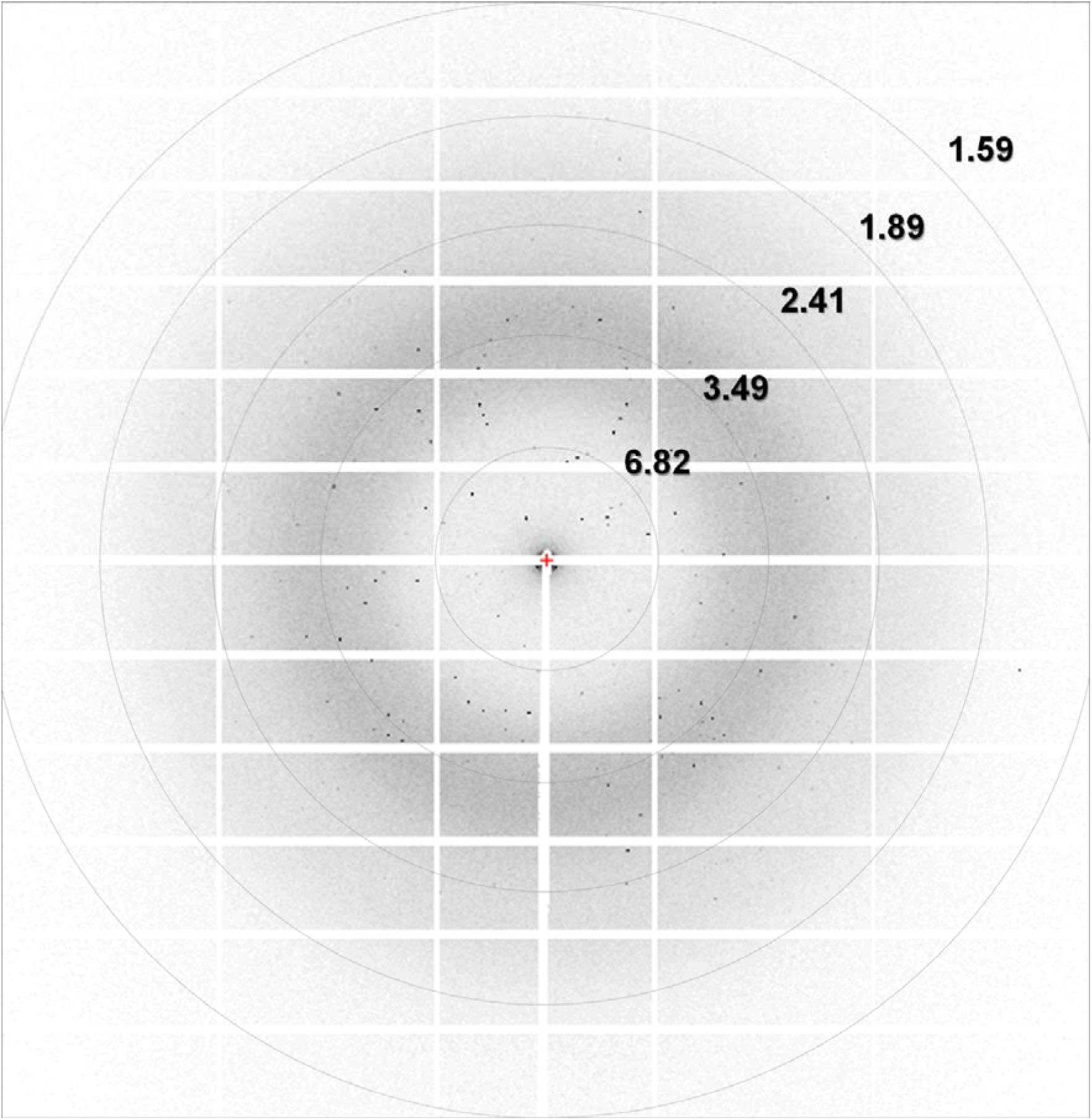
Diffraction pattern of glucose isomerase delivered in gelatin

**Supplementary Figure S3.**
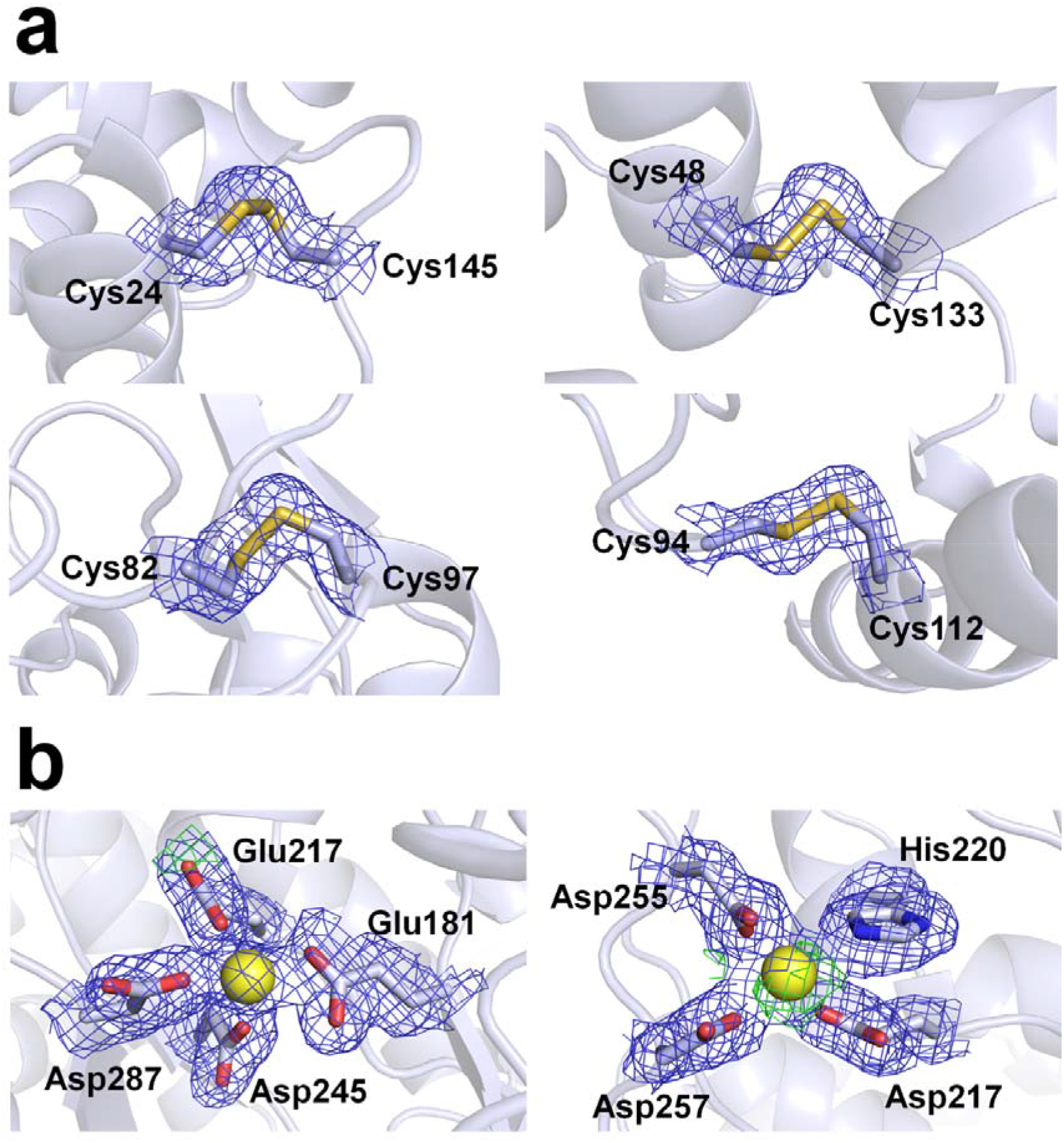
2Fo-Fc electron density map (grey, counted 1.5 σ) and Fo-Fc electron density map (green, counted 3 σ; red, counted - 3σ) of (a) disulfide bonds of lysozyme delivered in agarose and (b) metal binding sites of glucose isomerase delivered in gelatin.

## REFERENCES

1 Su, X. D. et al. Protein Crystallography from the Perspective of Technology Developments. Crystallogr Rev 21, 122–153 (2015).

2 Jaskolski, M., Dauter, Z. & Wlodawer, A. A brief history of macromolecular crystallography, illustrated by a family tree and its Nobel fruits. Febs J 281, 3985–4009 (2014).

3 Blundell, T. L. Protein crystallography and drug discovery: recollections of knowledge exchange between academia and industry. Iucrj 4, 308–321 (2017).

4 Holton, J. M. & Frankel, K. A. The minimum crystal size needed for a complete diffraction data set. Acta Crystallogr D 66, 393–408 (2010).

5 Helliwell, J. R. Protein Crystal Perfection and the Nature of Radiation-Damage. J Cryst Growth 90, 259–272 (1988).

6 Meents, A., Gutmann, S., Wagner, A. & Schulze-Briese, C. Origin and temperature dependence of radiation damage in biological samples at cryogenic temperatures. P Natl Acad Sci USA 107, 1094–1099 (2010).

7 Owen, R. L., Rudino-Pinera, E. & Garman, E. F. Experimental determination of the radiation dose limit for cryocooled protein crystals. P Natl Acad Sci USA 103, 4912–4917 (2006).

8 Chapman, H. N., Caleman, C. & Timneanu, N. Diffraction before destruction. Philos Trans R Soc Lond B Biol Sci 369, 20130313 (2014).

9 Nogly, P. et al. Lipidic cubic phase serial millisecond crystallography using synchrotron radiation. Iucrj 2, 168–176 (2015).

10 Weinert, T. et al. Serial millisecond crystallography for routine room-temperature structure determination at synchrotrons. Nat Commun 8, 542 (2017).

11 Schulz, E. C. et al. The hit-and-return system enables efficient time-resolved serial synchrotron crystallography. Nat Methods 15, 901–904 (2018).

12 Grunbein, M. L. & Nass Kovacs, G. Sample delivery for serial crystallography at free-electron lasers and synchrotrons. Acta Crystallogr D Struct Biol 75, 178–191 (2019).

13 DePonte, D. P. et al. Gas dynamic virtual nozzle for generation of microscopic droplet streams. J Phys D Appl Phys 41 (2008).

14 Weierstall, U. et al. Lipidic cubic phase injector facilitates membrane protein serial femtosecond crystallography. Nat. Commun. 5 (2014).

15 Hunter, M. S. et al. Fixed-target protein serial microcrystallography with an x-ray free electron laser. Sci Rep 4, 6026 (2014).

16 Lee, D. et al. Nylon mesh-based sample holder for fixed-target serial femtosecond crystallography. Sci Rep 9, 6971 (2019).

17 Sierra, R. G. et al. Nanoflow electrospinning serial femtosecond crystallography. Acta Crystallogr D 68, 1584–1587 (2012).

18 Monteiro, D. C. F. et al. A microfluidic flow-focusing device for low sample consumption serial synchrotron crystallography experiments in liquid flow. J Synchrotron Radiat 26, 406–412 (2019).

19 Nam, K. H. Sample Delivery Media for Serial Crystallography. Int J Mol Sci 20 (2019).

20 Boutet, S. et al. High-Resolution Protein Structure Determination by Serial Femtosecond Crystallography. Science 337, 362–364 (2012).

21 Park, J. et al. Polyacrylamide injection matrix for serial femtosecond crystallography. Sci Rep 9, 2525 (2019).

22 Stellato, F. et al. Room-temperature macromolecular serial crystallography using synchrotron radiation. Iucrj 1, 204–212 (2014).

23 Lopez-Jaramillo, F. J., Garcia-Ruiz, J. M., Gavira, J. A. & Otalora, F. Crystallization and cryocrystallography inside X-ray capillaries. J Appl Crystallogr 34, 365–370 (2001).

24 Conrad, C. E. et al. A novel inert crystal delivery medium for serial femtosecond crystallography. Iucrj 2, 421–430 (2015).

25 Park, S. Y., Ha, S. C. & Kim, Y. G. The Protein Crystallography Beamlines at the Pohang Light Source II. Biodesign 5, 30–34 (2017).

26 Park, S. Y. & Nam, K. H. Sample delivery using viscous media, syringe, and syringe pump for serial crystallography. J Synchrotron Radiat (2019).

27 Barty, A. et al. Cheetah: software for high-throughput reduction and analysis of serial femtosecond X-ray diffraction data. J Appl Crystallogr 47, 1118–1131 (2014).

28 White, T. A. et al. Recent developments in CrystFEL. J Appl Crystallogr 49, 680–689 (2016).

29 Adams, P. D. et al. PHENIX: a comprehensive Python-based system for macromolecular structure solution. Acta Crystallogr D Biol Crystallogr 66, 213–221 (2010).

30 Bae, J. E., Hwang, K. Y. & Nam, K. H. Structural analysis of substrate recognition by glucose isomerase in Mn(2+) binding mode at M2 site in S. rubiginosus. Biochem Biophys Res Commun 503, 770–775 (2018).

31 Emsley, P. & Cowtan, K. Coot: model-building tools for molecular graphics. Acta Crystallogr D Biol Crystallogr 60, 2126–2132 (2004).

32 Adams, P. D. et al. PHENIX: a comprehensive Python-based system for macromolecular structure solution. Acta Crystallogr D 66, 213–221 (2010).

33 Williams, C. J. et al. MolProbity: More and better reference data for improved all-atom structure validation. Protein Sci 27, 293–315 (2018).

